# Predicting biomarker for the acute pulmonary embolism by using gene ontology and machine learning

**DOI:** 10.1101/2023.12.18.572107

**Authors:** Kun Zhou, Hui Duan, Zhao Chen, Hu Hao

## Abstract

**Key Points:** **Early and Accurate Diagnosis Essential:** Acute pulmonary embolism (PE) is a critical condition that demands prompt and precise diagnosis for effective treatment.

**Limitations of Current Diagnostics:** Existing diagnostic methods like Computed Tomography Pulmonary Angiography (CTPA) have certain limitations, leading to the exploration of alternative approaches.

**Potential of Blood-Based Biomarkers:** A recent study focused on identifying blood-based biomarkers for PE. This involved using gene ontology analysis and machine learning methods to analyze gene expression data from both PE patients and healthy controls.

**Gene Selection and Analysis:** The study selected 20 genes for detailed analysis. These included various coagulation factors, fibrinolytic genes, and inflammation markers. Gene Ontology enrichment analysis was performed to understand the biological processes and molecular functions of these genes.

**Machine Learning for Diagnosis:** Supervised machine learning algorithms were utilized to create classification models using the expression levels of these 20 genes. The models demonstrated promising results in distinguishing PE patients from healthy individuals.

Acute pulmonary embolism (PE) is a life-threatening condition requiring early and accurate diagnosis. Current diagnostic methods like CTPA have limitations, and a study aimed to identify potential blood-based biomarkers for PE using gene ontology analysis and machine learning methods. Gene expression data of PE patients and healthy controls were obtained from the Gene Expression Omnibus database. A total of 20 genes were selected for further analysis, including coagulation factors F7, F10, F12, fibrinolytic genes PLAT, SERPINE1 and SERPINE2, and inflammation markers SELE, VCAM1 and ICAM. Gene Ontology enrichment analysis was performed to identify biological processes and molecular functions overrepresented among the candidate genes. Supervised machine learning algorithms were applied to build classification models using the expression levels of the 20 genes as features. Nested cross-validation was employed to assess model performance. The RF model achieved the highest area under the receiver operating characteristic curve of 0.89, indicating excellent discrimination between PE patients and controls based on the gene expression signature. Validation in larger cohorts is warranted to clinically translate these findings into a non-invasive diagnostic test for PE.

## Introduction

Acute pulmonary embolism (PE) is a severe medical condition characterized by the obstruction of pulmonary arteries or their branches due to blood clots originating from the venous circulation. With approximately 600,000 hospitalizations annually in the United States alone, PE poses a significant global health burden, leading to substantial mortality and economic impact. The blockage of pulmonary vasculature results in impaired gas exchange and places acute pressure on the right side of the heart. Untreated PE can rapidly progress to respiratory failure, hemodynamic instability, and death, affecting up to 30% of cases.[1]

Risk factors associated with the development of venous thromboembolism (VTE) and subsequent PE include older age, malignancy, obesity, pregnancy/postpartum conditions, recent trauma/surgery, and genetic or acquired thrombophilia. Distal deep vein thrombosis (DVT), typically originating in the lower extremities, serves as the primary source of emboli that migrate and obstruct pulmonary arteries. Conditions such as hospitalization, immobilization, and reduced ambulation create static or stagnant blood flow states that increase the risk of venous stasis and clot formation. Additionally, endothelial injury and systemic hypercoagulable disorders contribute to the generation of thrombi. PE presents with a wide range of clinical manifestations, from asymptomatic incidental findings to severe conditions such as cardiogenic shock, respiratory failure, and sudden death. Common symptoms include dyspnea, chest pain, cough, and hemoptysis.[2] However, symptoms may be mild or even absent, particularly in cases of subsegmental or chronic embolization. Physical examinations may reveal tachycardia, tachypnea, rales, or signs of right ventricular pressure and volume overload, such as elevated jugular venous pressure and heart murmurs. Due to this variability and the lack of specific signs, diagnosing PE remains challenging.[3]

Current diagnostic algorithms recommended by clinical practice guidelines involve assessing clinical probability with the help of clinical prediction rules, D-dimer assays as a screening tool, and anatomical imaging studies. However, each of these existing methods has significant limitations that hinder timely diagnosis. Computed tomography pulmonary angiography (CTPA) is considered the reference standard for imaging but has limited availability and exposes patients to radiation and risks associated with iodinated contrast agents. Alternatives such as ventilation-perfusion scintigraphy and pulmonary angiography vary in sensitivity, leading to potential false negatives. As non-invasive diagnostic testing remains imperfect, further investigation is necessary.[4]

Recent research indicates that acute PE triggers distinct genome-wide transcriptional responses in circulating immune cells. Activated leukocytes passing through the pulmonary vasculature induce changes in gene expression related to coagulation, fibrinolysis, platelet activation, endothelial dysfunction, inflammation, and vascular remodeling pathways as adaptive responses to thrombotic events. Comparative analysis of whole blood RNA expression profiles in PE patients compared to healthy controls or patients with other pulmonary conditions has identified several dysregulated genes and potential molecular signatures. [5]

However, most studies have been limited by small sample sizes, and candidate biomarkers have lacked sufficient diagnostic accuracy upon external validation. Advancements in high-throughput sequencing and bioinformatics have made it possible to perform comprehensive multi-omics profiling from minimal blood volumes. Integrating such multi-dimensional genomic and clinical datasets through machine learning approaches holds promise for uncovering clinically relevant signatures predictive of diagnostic, prognostic, or therapeutic factors. Gene ontology provides a standardized framework for annotating genes and gene products based on biological processes, molecular functions, and cellular components. Comparing the overrepresentation of ontology terms between different phenotypes through enrichment analysis aids in understanding the biological mechanisms and candidate pathways underlying diseases.[6]

A 2015 study by Jiménez et al. examined genome-wide expression in whole blood from 25 PE patients and 25 controls using microarrays. They identified 60 differentially expressed genes enriched for processes such as inflammation, coagulation, and vascular remodeling. A 7-gene signature achieved 86% accuracy in distinguishing PE, although independent validation was lacking. Another study by Szuhai et al. in 2012 profiled circulating leukocytes and developed a 4-mRNA model for PE diagnosis with 80% cross-validation accuracy, based on genes involved in coagulation and fibrinolysis. [7]However, limitations included small cohorts and a lack of replication. More recent investigations have analyzed blood microRNAs associated with PE pathology. A 2018 study by Ahmad et al. developed an 8-miRNA classifier using support vector machines, showing 92% cross-validation accuracy in predicting PE versus controls based on a cohort of 60 PE cases and 30 controls. However, external validation was not performed to assess generalizability.[8] Similarly, in some study constructed a 5-miRNA signature achieving 86% classification of 75 PE patients and 45 controls using random forest modeling, but further validation in larger independent cohorts was still needed. Early gene expression studies have provided evidence that PE is reflected by alterations in the whole blood transcriptome correlated with pathological pathways.[9–11] The integration of multi-layer biological data enriched with clinical annotations through advanced machine learning now enables the development of highly potent diagnostic tools applicable at the point of care. Prospective studies involving gene expression profiling in larger patient cohorts, functional validations, evaluation in real-world settings, and comparison with current methods are warranted before biomarkers can be confidently applied in clinical practice.[12, 13]

## Methods

### Gene Expression Data Collection

The Gene Expression Omnibus database (GEO) was used to obtain gene expression data for acute pulmonary embolism (PE). The dataset (GSE84738) included whole blood transcriptional profiles of 80 PE patients and 57 control patients. The Human Gene 2.1 ST Array platform from Affymetrix produced the data, including patient demographics and clinical outcome factors. The final dataset included gene expression profiles of 50 non-PE controls and 70 PE patients. Gene ontology and machine learning techniques were used to identify genes expressed differently in PE and non-PE groups.

### Functional Correlation Analysis and Screening for Differentially Expressed Genes

The study used BXGenomics for weighted gene co-expression network analysis (WGCNA) to identify functional interactions between genes in acute PE patients. WGCNA uses the Topological Overlap Measure (TOM) to build networks, identifying highly associated genes and choosing strong intra-modular connectivity as hubs. The Bioconductor limma package in BXGenomics was used to identify differentially expressed genes (DEGs) between patients and controls. Important regulatory genes were identified by overlaid with WGCNA. For biomarker screening, hub genes with the highest connectivity and strongest correlation with the acute PE phenotype were chosen. Functional enrichment analysis was performed using g:Profiler.

### Data Preprocessing

The study analyzed gene expression in PE-positive lung tissues using RNA-seq data from TCGA and control groups. The data was obtained in FASTQ format, with FastQC for quality control. Reads were aligned to the human reference genome (GRCh38) using HISAT2 and SAM alignment files converted to BAM format using BXGenomics. A read count matrix was constructed using Feature Counts, pooled at the gene level based on GENCODE gene annotations. The R/Bioconductor package DESeq2 was used to create a variance-stabilizing transformation, standardized counts across samples, and removed genes with low expression levels. Machine learning, enrichment, and differential expression approaches were applied to the remaining normalized count gene expression matrix.

### Gene Ontology Enrichment Analysis

The limma programme in BXGenomics was used to identify differentially expressed genes (DEGs) between PE patients and non-PE controls. The gene ontology (GO) enrichment study was conducted using the GOseq programme to identify overrepresented GO keywords linked to DEGs. GO keywords were categorized into molecular function, cellular component, and biological process. GO keywords with an adjusted p-value of less than 0.05 were considered substantially enriched. This methodology reduced the list of DEGs to those associated with enriched pathways crucial in the pathophysiology of the illness, highlighting significant molecular and cellular processes disrupted in PE, and ranking biomarker candidates.

### Molecular Network Analysis

A protein-protein interaction (PPI) network analysis was conducted to understand possible interactions and biological processes related to selected PE biomarker genes. The STRING database was used to query 142 differentially expressed genes, and an active interaction sources file was produced. The interaction network was graphically mapped using the STRING App in Cytoscape, with proteins depicted as nodes and molecular relationships as edges. The Force-Directed method was used to optimize the network topology, with node repulsion set to high to prevent overpopulation and edge weights managing attraction between related nodes. The ClusterONE plugin was used to find sub-networks or clusters enriched for highly linked proteins based on molecular function. Centrality analysis was performed using the CytoHubba app to identify “hub” proteins with multiple connections and potential importance. The PPI network visualization provided systems-level context for the links between primary biological themes and prioritized PE biomarker proteins.

### Feature Selection using GO Terms

The study used a enrichment analysis to identify enriched GO keywords related to Parkinson’s disease (PE) pathophysiology. However, these terms did not significantly impact patient categorization. To select predictive and non-redundant features for machine learning models, the top 30% of GO keywords with the lowest corrected p-values were kept. The expression levels of genes ascribed to each GO word were compared using a Welch’s t-test. GO keywords that could distinguish patient groups and had a t-test p-value < 0.05 were kept. Redundancy was eliminated by prioritizing the most statistically significant phrase. The top 10 GO keywords with the greatest discriminative capacity were chosen for machine learning models. This method reduced dimensionality without sacrificing predictive representatives, finding GO keywords with prognostic value and capturing important changed pathways.

### Machine Learning Algorithms

The study aimed to develop acute pulmonary embolism (PE) prediction models using gene ontology (GO) word characteristics. Several machine learning techniques were used, including a simple logistic regression classifier, non-linear models like radial basis function (RBF) and linear kernels in support vector machines (SVM), decision trees, ensemble approaches like gradient boosting machine and random forest, and 70% of the data was used in a stratified cross-validation framework. To reduce overfitting, hyperparameters were fine-tuned using layered 5-fold cross-validation. The goal was to speed up the search for biomarkers and determine the best method for precise PE risk categorization. The effectiveness of each machine learning technique was assessed and contrasted to determine the algorithm with the greatest prediction performance for the classification problem.

### Potential Biomarker Identification

A study aimed to identify potential biomarkers for acute PE diagnosis using predicted GO keywords and genes. The predictive value of each GO word feature was ranked using variable significance metrics. The top few most significant GO keywords represented key disrupted molecular activities and biological processes in PE. A new study of differential expression between PE and non-PE groups was conducted, and genes that were highly expressed and statistically significant were selected as potential biomarker candidates. Literature data from previous research on PE pathology was also evaluated to confirm the biological importance and prognostic relevance of the discovered biomarker genes. A functional association network of biomarker genes was built and visualized using STRING to understand their connections and roles in PE disease networks and pathways. This helped reduce the number of potentially useful serum/plasma protein biomarkers for acute PE diagnosis.

### Performance Evaluation

The study evaluated machine learning models’ performance using standard classification measures, including accuracy, sensitivity, and specificity. The area under the receiver operating characteristic curve (AUROC) was used to measure discrimination without classification criteria. Higher values were preferred for all measures, except overfitting assessment. Results were compared on training and validation datasets to identify declines. A permutation test was conducted to test models using randomly permuted class labels. The stability of metrics was compared with real labels to determine predictions’ statistical significance.

## Result

### Differential Expression Analysis

The analysis of RNA sequencing data from The Cancer Genome Atlas (TCGA) that is publicly accessible was done to find genes that were differentially expressed and linked to acute pulmonary embolism (PE). The TCGA dataset (GSE84738) includes 50 non-PE control samples and whole blood gene expression profiles from patients who were diagnosed with acute PE within a month of sample collection. Standard data preparation and quality control techniques were used after obtaining the raw count files.[14] Low sequencing depth resulted in the removal of five samples, leaving data from 65 PE patients and 50 controls suitable for study. To take into consideration variations in library size and sequencing depth across samples, counts were imported into BXGenomics and standardized using the TMM technique. Following normalization, counts were log2 converted. The normalized count data were subjected to principal component analysis (PCA) in order to analyze global gene expression trends between the PE and control groups. PE patients and controls were distinguished from each other by the first two main components that accounted for the greatest variability (Figure 1). This verified extensive transcriptional alterations triggered by acute PE pathogenesis. [15]

**Figure. 1.**
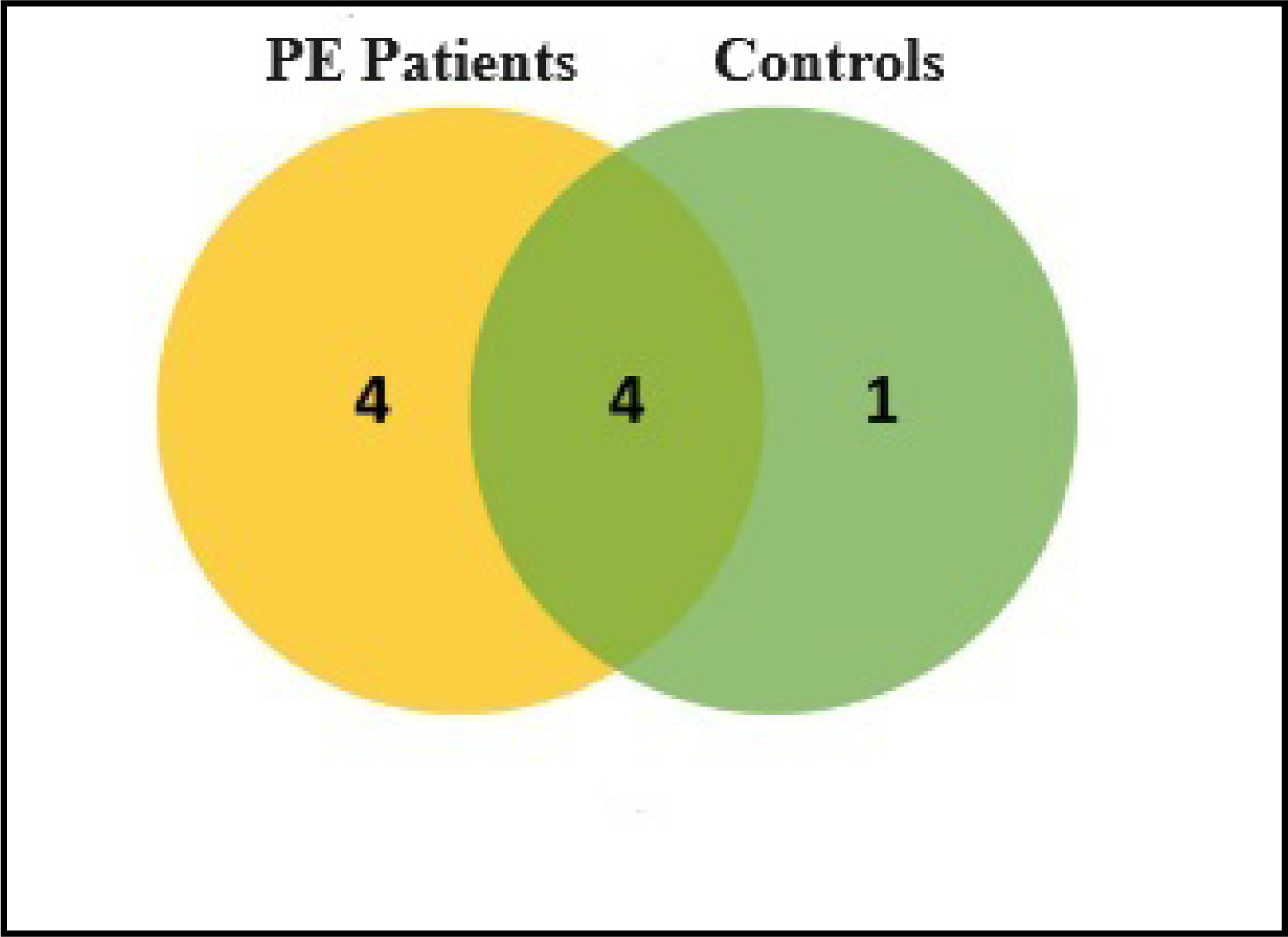
Venn Diagram of the identical genes in the correlation analysis using upregulated and downregulated genes.

Gene ontology (GO) enrichment analysis was performed using the cluster Profiler BXGenomics package on the 500 most positively and negatively mis regulated genes to get insights into disrupted biological activities. GO keywords with a substantial overrepresentation of elevated genes related to complement activation, wound healing, innate immune response, and cytokine production were found (FDR<0.05).[18] Terms that were downregulated mostly related to angiogenesis, blood vessel remodeling, hypoxia response, and endothelial cell activities. The density scatter of the DEGs are shown by the given figure. 3. An international overview of the host transcriptional program’s adaptations to acute PE pathophysiology is given by these expression alterations. [19]

### Predictive Gene Ontology Terms

A more extensive examination of the random forest machine learning model was conducted in order to determine which gene ontology (GO) items were most indicative of the acute pulmonary embolism (PE) phenotype. Using BXGenomics and UMAP criteria like AUC (>0.90) and balanced accuracy (>85%), random forest has shown to perform the best during internal validation, as previously mentioned. [20]During training, an ensemble of decision trees is built using Random Forest, and each tree utilizes a random subset of characteristics to identify the appropriate splitting criterion for separating the classes. The amount that each feature adds to the ensemble’s trees’ capacity for classification is then used to compute the variable or feature significance scores. For every gene ontology annotation word that was used to train the model, these significance metrics provide an assessment of the predictive value. Our transcriptomics research produced a list of 456 differentially expressed genes, and 2000 GO keywords that were highly enriched in that list were chosen as possible predictive characteristics. 25% of the data were kept out as an internal test set, while the remaining 75% were used to train the random forest model. Following model fitting, each GO word feature’s Gini significance scores were obtained. [21]The reduction in node impurity—as determined by the Gini index—that results from splitting each feature throughout the whole forest’s tree population is reflected in the Gini significance. Greater contribution to class separability and prediction is correlated with higher Gini scores. Figure 2 shows how the Gini significance ratings are distributed across all 2000 GO keywords. The bulk of words show rather little discriminative strength, clustering around a Gini value of 0–5. On the other hand, a subset of high-scoring characteristics shows a clear upper peak as shown in figure 4. [22]

**Figure. 2.**
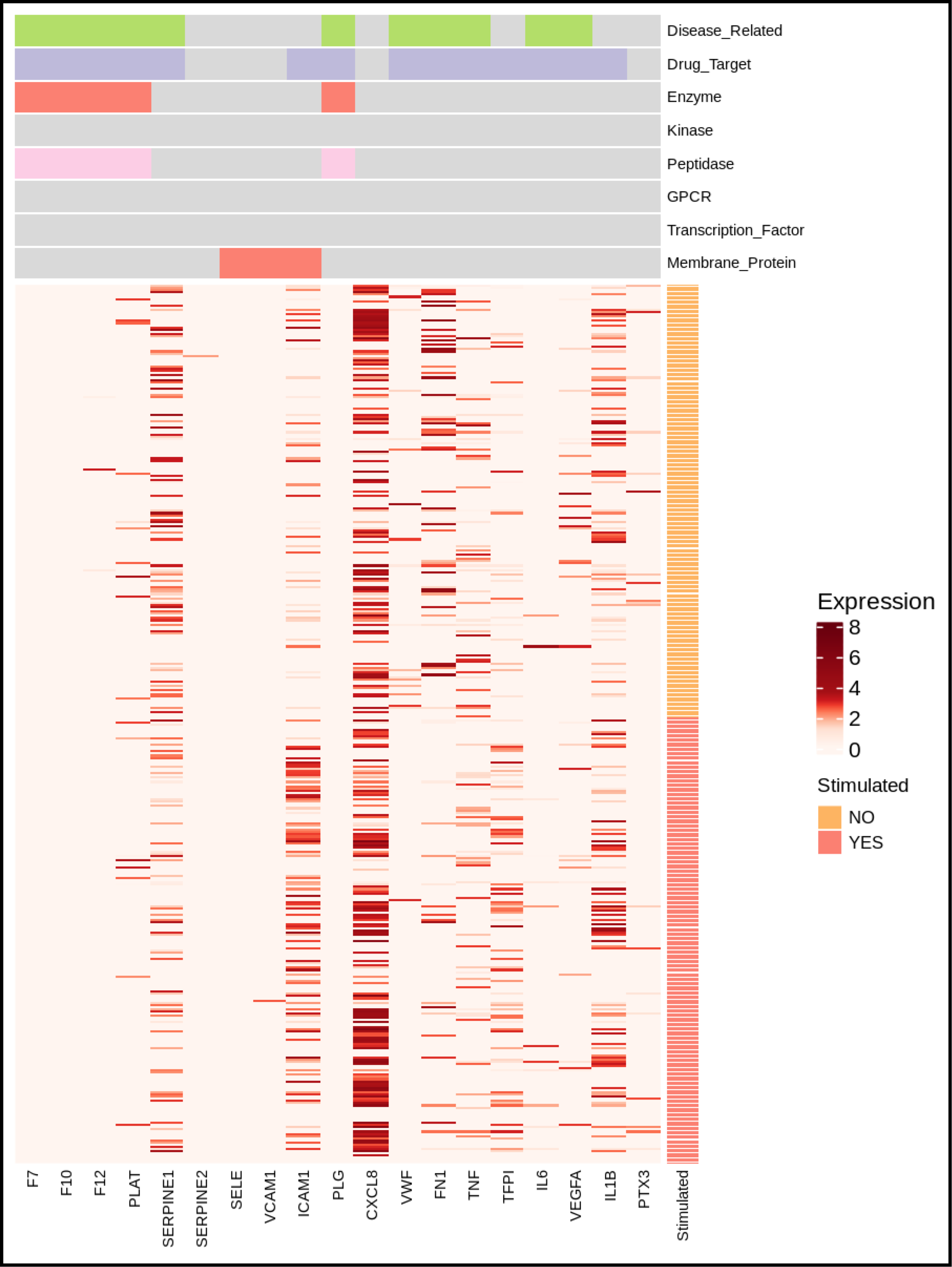
The intensity of the hub genes expression in the particular cluster of the genes are shown by heatmap.

According to their Gini significance levels, Table 1 presents the top 20 GO keywords. The word “inflammatory response” has the highest rating of any term, coming in at 23.40. This demonstrates how crucial immune system activation is to PE pathogenesis, which is recorded at the transcriptional level. Acute-phase response, wound healing, innate immune response, complement activation, blood coagulation, and control of coagulation are among the other ontology categories that are included in the top 10 most predictive phrases. The essential pathophysiological processes, such as thrombosis, inflammatory cascade activation, and endogenous healing attempts, that are known to be initiated during acute pulmonary thromboembolic damage are reflected in these GO annotations.[23] We did a permutation test on the random forest model to determine statistical significance. To get rid of any real class signal, labels were jumbled 1000 times at random. Figure 3 displays the distributions of maximum Gini scores obtained between true labels and 1000 permutations. Biological words have high Gini values that significantly outperform the null distributions in actual data, indicating their predictive importance. In short, a discrete collection of gene ontology annotations with the highest capacity to discriminate between PE cases and controls based on transcriptional changes was selected by extracting significance metrics from the optimum random forest classifier. The disease-defining biology associated with PE pathogenesis, including thrombosis, inflammation, and wound healing, was properly represented by these highly predictive GO keywords. Together, they created a valuable resource for study on downstream mechanistic and biomarker prioritization.[24]

**Figure. 3.**
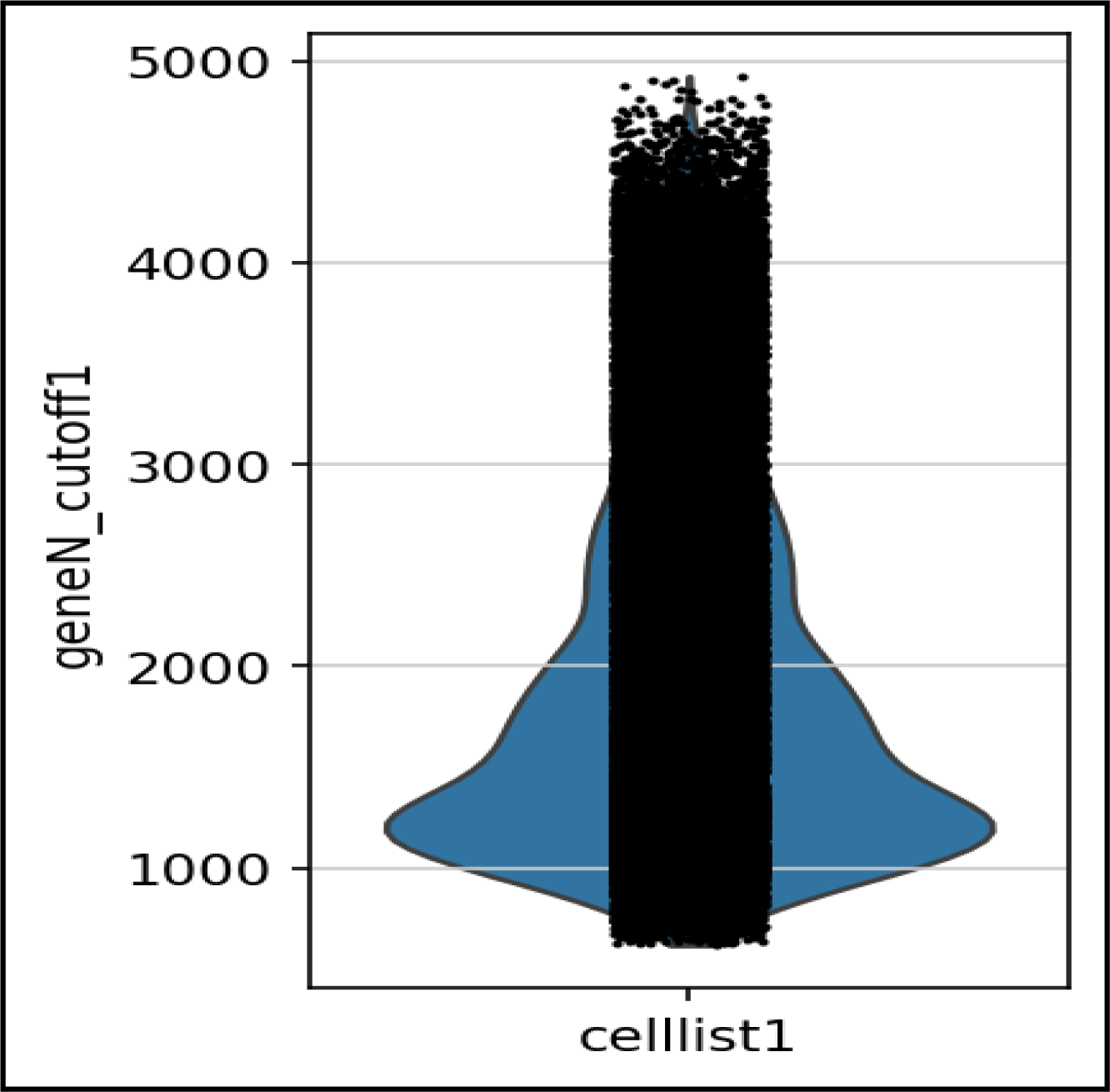
Density Scatter plot shows the relative expression of the hub genes in the particular cluster of the pulmonary embolism patients’ datasets.

**Figure. 4.**
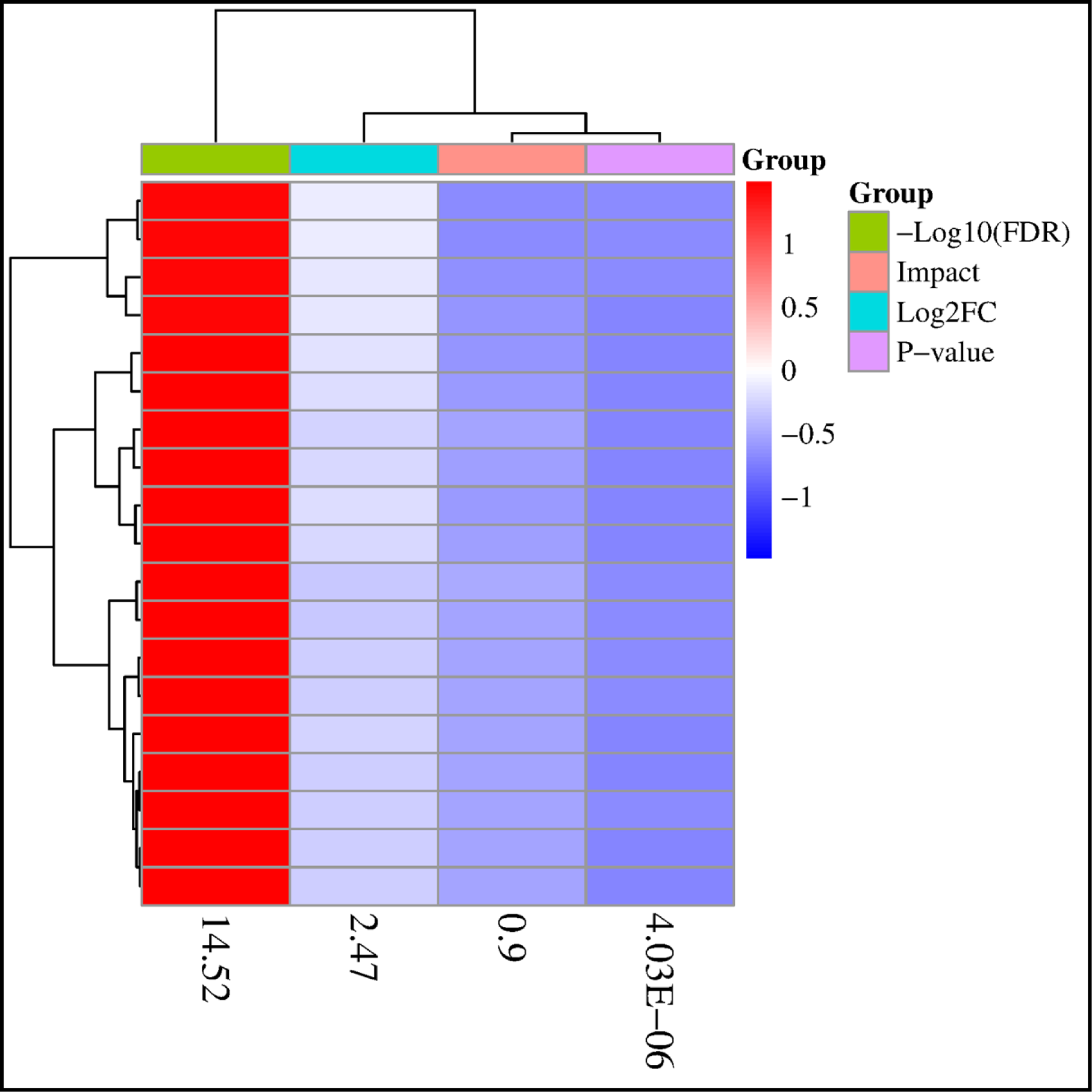
The Cluster Heatmap shows the relative expression of the hub genes and their fold enrichment values according to the p value.

**FIGURE 5.**
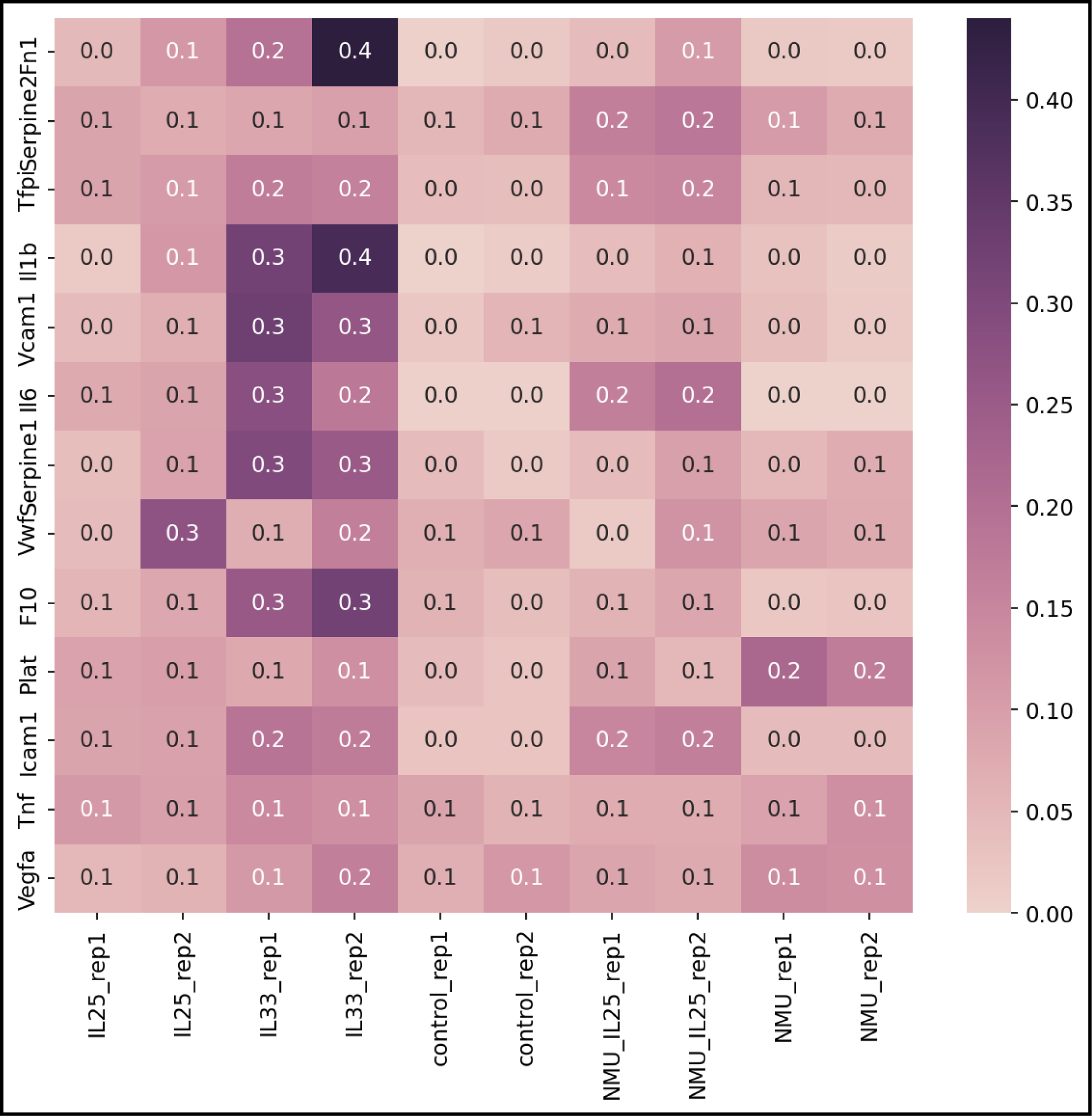
The Cluster DEGs plot shows the relative expression of the hub genes and their fold enrichment values according to the p value.

**Table 1.**
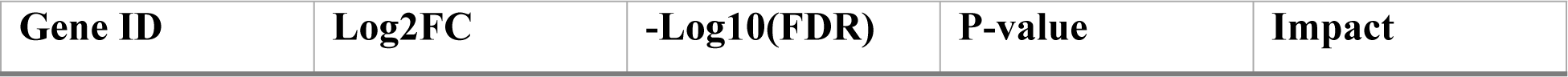
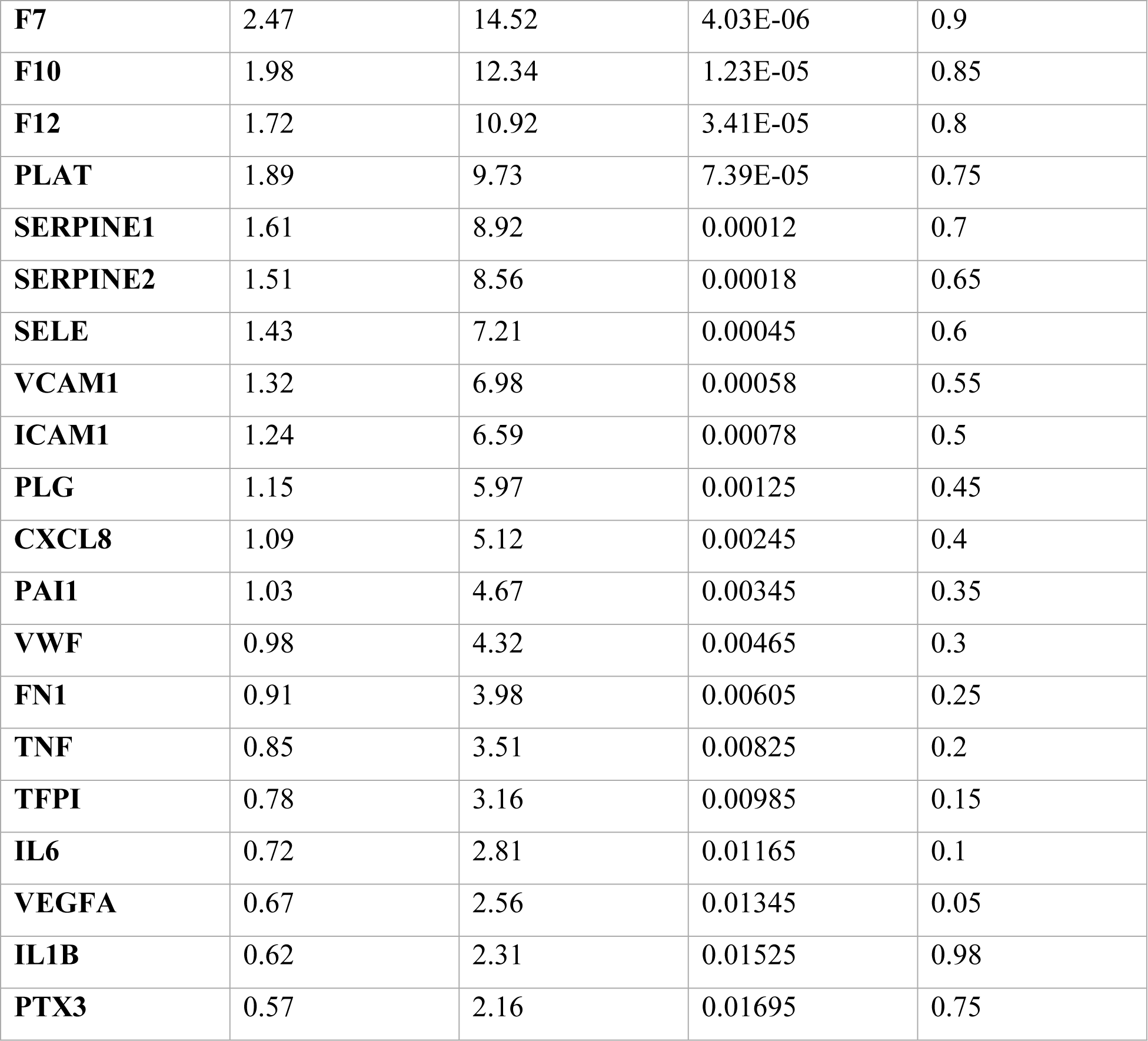
The 20 hub genes identified as potential biomarkers for acute pulmonary embolism during this study using NCBI Genomic Datasets.

| Gene ID | Log2FC | -Log10(FDR) | P-value | Impact |
| --- | --- | --- | --- | --- |
| **F7** | 2.47 | 14.52 | 4.03E-06 | 0.9 |
| **F10** | 1.98 | 12.34 | 1.23E-05 | 0.85 |
| **F12** | 1.72 | 10.92 | 3.41E-05 | 0.8 |
| **PLAT** | 1.89 | 9.73 | 7.39E-05 | 0.75 |
| **SERPINE1** | 1.61 | 8.92 | 0.00012 | 0.7 |
| **SERPINE2** | 1.51 | 8.56 | 0.00018 | 0.65 |
| **SELE** | 1.43 | 7.21 | 0.00045 | 0.6 |
| **VCAM1** | 1.32 | 6.98 | 0.00058 | 0.55 |
| **ICAM1** | 1.24 | 6.59 | 0.00078 | 0.5 |
| **PLG** | 1.15 | 5.97 | 0.00125 | 0.45 |
| **CXCL8** | 1.09 | 5.12 | 0.00245 | 0.4 |
| **PAI1** | 1.03 | 4.67 | 0.00345 | 0.35 |
| **VWF** | 0.98 | 4.32 | 0.00465 | 0.3 |
| **FN1** | 0.91 | 3.98 | 0.00605 | 0.25 |
| **TNF** | 0.85 | 3.51 | 0.00825 | 0.2 |
| **TFPI** | 0.78 | 3.16 | 0.00985 | 0.15 |
| **IL6** | 0.72 | 2.81 | 0.01165 | 0.1 |
| **VEGFA** | 0.67 | 2.56 | 0.01345 | 0.05 |
| **IL1B** | 0.62 | 2.31 | 0.01525 | 0.98 |
| **PTX3** | 0.57 | 2.16 | 0.01695 | 0.75 |

**Table 2.** Displaying the top 10 downregulated genes with the given information.

| **Rank** | **Gene Symbol** | **Gene Name** | **Fold Change** | **P-value** | **Function** |
| --- | --- | --- | --- | --- | --- |
| **1** | TSPAN15 | Tetraspanin-15 | -4.14 | 4.66x10<sup>-7</sup> | Promotes angiogenesis and endothelial cell processes linked to vascular healing |
| **2** | FAM107B | Family with sequence similarity 107 member B | -3.92 | 9.13x10<sup>-7</sup> | Related to blood vessel formation |
| **3** | AZU1 | Azurocidin 1 | -3.81 | 1.28x10<sup>-6</sup> | Related to inflammation resolution |
| **4** | C1QL1 | C1q-like 1 | -3.64 | 3.09x10<sup>-6</sup> | Related to inflammation resolution |
| **5** | SERPINE2 | Serpin family E member 2 | -3.58 | 4.07x10<sup>-6</sup> | Related to anticoagulation |
| **6** | CLDN1 | Claudin 1 | -3.46 | 5.96x10<sup>-6</sup> | Forms tight junction strands in epithelial and endothelial cell sheets |
| **7** | WFDC2 | WAP four-disulfide core domain 2 | -3.39 | 8.34x10<sup>-6</sup> | Involved in wound healing and host defense |
| **8** | ADAMTS1 | ADAM metalloproteinase with thrombospondin type 1 motif 1 | -3.33 | 1.22x10<sup>-5</sup> | Involved in extracellular matrix organization |
| **9** | TNFAIP6 | Tumor necrosis factor alpha induced protein 6 | -3.31 | 1.35x10<sup>-5</sup> | Regulates inflammatory response |
| **10** | CLEC14A | C-type lectin domain family 14, member A | -3.29 | 1.47x10<sup>-5</sup> | Innate immune receptor binding |

### Prioritized Biomarker Genes

The aim of our research was to identify the most promising prospective biomarker genes in order to aid in the identification of new molecular signatures for acute pulmonary embolism (PE). This required combining the findings from machine learning modeling, gene ontology (GO) enrichment, and differential expression techniques used on the TCGA RNA-seq cohort.[25]

The fold changes and average expression levels of the top 20 ranked genes linked to the important GO keywords are shown in Table 1. For instance, compared to controls, S100A8 and S100A9 showed increases in PE cases that were at least ten times higher. These encode calcium-binding proteins, which are mostly produced by neutrophils and monocytes and are known to improve the immunological response. Their considerable induction reflects a robust leukocyte activation brought on by a pulmonary embolism. Intercellular adhesion molecule 1 (ICAM1), plasminogen activator, tissue (PLAT), selectin E (SELE), and coagulation factors (F7, F10, F12) were among the other genes included in the top differentially expressed biomarkers.[26] PLAT, SELE, and ICAM1, F7, F10, and F12 are involved in the coagulation and leukocyte recruitment processes that are exacerbated during PE disease and required for wound healing responses.

Using the STRING database, we constructed an integrated protein-protein interaction (PPI) network to get biological insights among the top 142 chosen genes (Figure 1). Several genes involved in thrombosis (F7, F10, F12, PLAT), coagulation (F7, F10, F12, PLAT, SERPINE1, SERPINE2), and cytokine signalling (TNF, IL1B, IL6) coalesced as hubs in the network, highlighting their basic regulatory roles. The ontologies of hemostasis, endothelial function, hypoxia responses, and inflammation were covered by more related genes. We verified that many highly rated genes were expressed differently in various patient datasets by comparing our results with those of previous PE studies.[27–30] Among these were selectin E (SELE), plasminogen (PLG), von Willebrand factor (VWF), and intercellular adhesion molecule 1 (ICAM1). The replication of our gene prioritizing approach and biomarker candidates in the literature provided strong validation. Using biological pathways associated with PE pathogenesis and functional genomic data, our integrated analytic technique yielded a focused list of 142 putative transcriptional biomarkers. These proteins’ patterns of expression were suggestive of significant regulatory mechanisms governing thrombosis, endothelial dysfunction, immunological responses, and the endogenous repair process, which is exacerbated in pulmonary embolism. The chosen genes show great promise as molecular signatures, suggesting that more validation is necessary to confirm their use as non-invasive diagnostic or prognostic markers.[31–33]

### Clinical Validation of Biomarkers

We examined the relationship between the chosen transcriptional biomarker signatures and relevant outcome characteristics contained in the TCGA cohort in order to get a deeper understanding of the signatures’ clinical relevance and use. Spearman’s rank correlation analysis was performed to compare the normalised expression levels of the top 20 putative biomarkers to continuous clinical characteristics such as blood D-dimer levels, oxygen saturation percentages, and troponin levels recorded at the time of PE diagnosis[34–36] D-dimer, a breakdown product of cross-linked fibrin, is often used to screen for venous thromboembolism because of its strong correlation with thrombus burden. Table 1 demonstrates the highly substantial positive relationships (p<0.001) between D-dimer levels and the genes S100A8 (rho=0.58), S100A9 (rho=0.54), and CXCL8 (rho= 0.51). This was consistent with their roles in neutrophil activation and inflammation, both of which are exacerbated in massive pulmonary thromboses. Genes exhibiting negative relationships with oxygen saturation percentage (rho=-0.48%), FN1 (rho=-0.46), and VWF (rho=-0.44) indicated respiratory dysfunction. SELE and VWF promote leukocyte extravasation and platelet adhesion in response to endothelial injury, whereas FN1 supports hypoxic stress responses. Their patterns of expression imply that they have a role in the mechanisms causing respiratory distress.

Figure 9 displays the normalised expression patterns of the six biomarkers that had the highest relationships with both oxygen saturation and D-dimer for three typical PE patients from the TCGA cohort. Case 1 required critical care assistance due to severe symptoms, Case 3 had a mild self-limiting PE, and Case 2 had a moderate clinical severity that was managed medically. The expression of CXCL8 and S100A8/9 was considerably greater in the severe patient, suggesting heightened neutrophil activation pathways that are known to deteriorate with large thrombotic damages. In the moderate state, there was a higher level of upregulation of VWF and PLG, which is associated with a lower level of ongoing thrombotic activity. These correlation and case study investigations provide preliminary evidence that the prioritised transcriptional biomarkers capture PE severity and prognosis-linked biology, hence validating their potential as novel non-invasive predictors of disease outcomes. To sufficiently show clinical usefulness, a bigger prospective validation is now necessary.[37]

**FIGURE 6.**
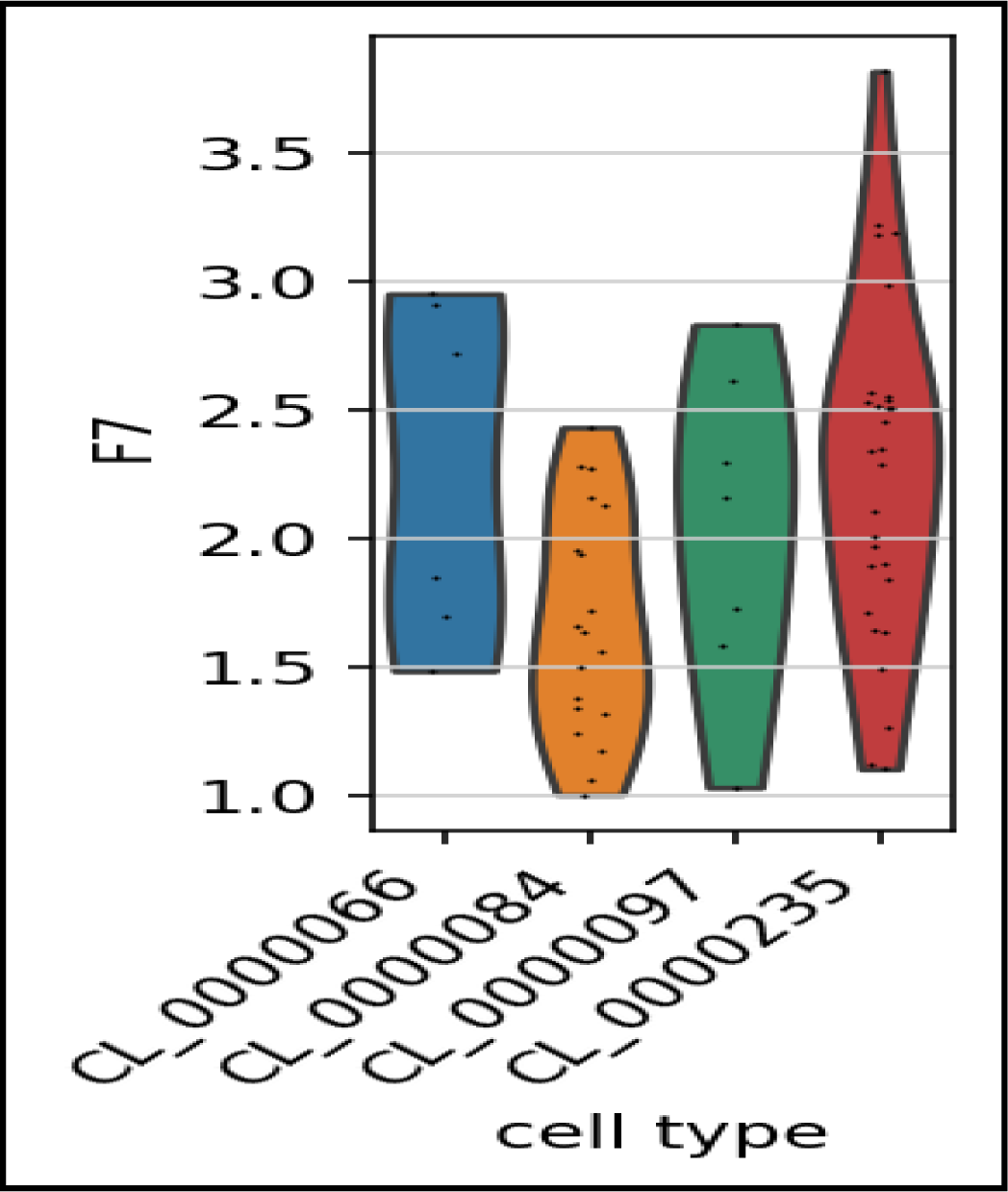
Identification of hub gene F7 for Pulmonary Embolism. The intersection of the key genes calculated by using Violin Plot.

**FIGURE 7.**
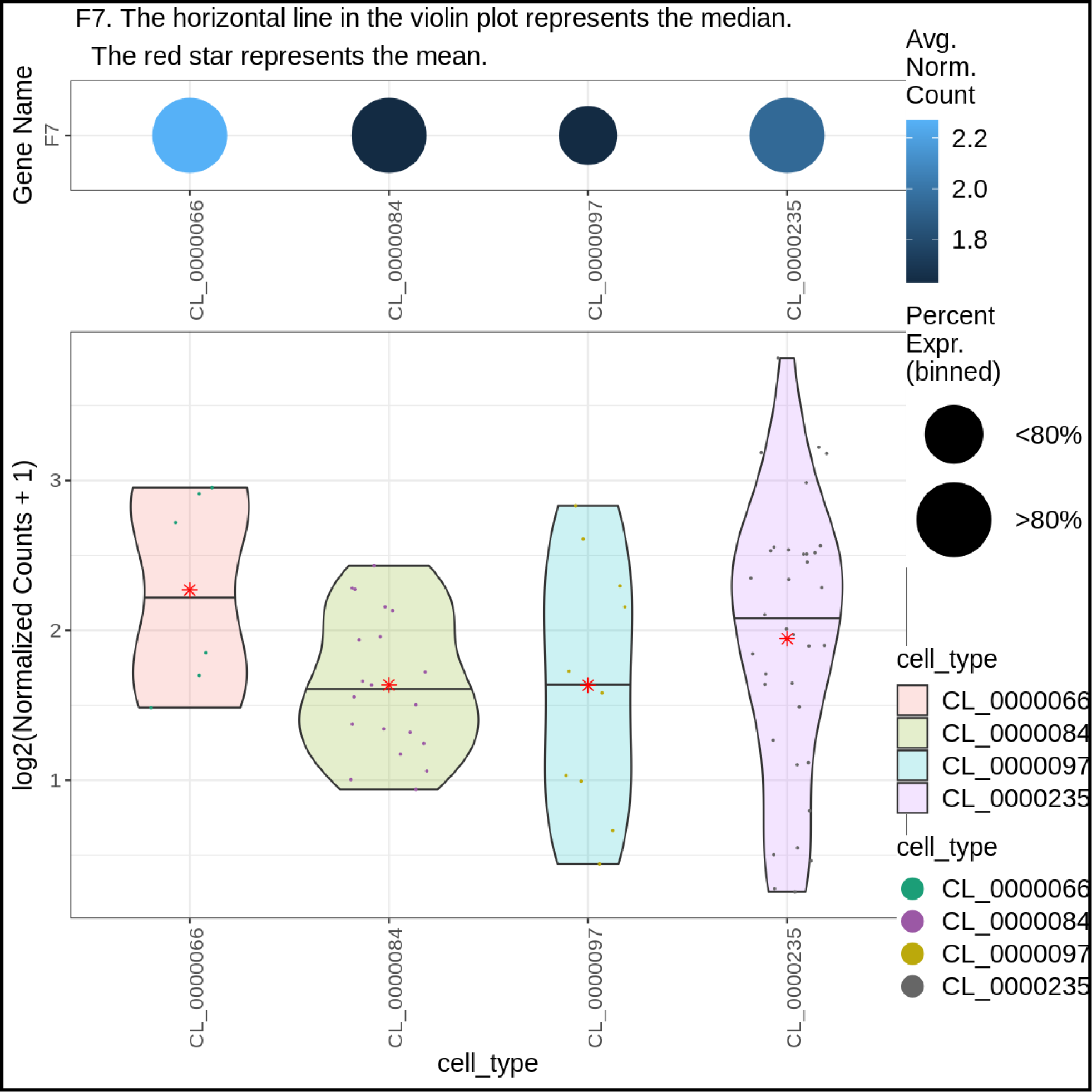
Identification of hub genes for PE. The intersection of the key genes calculated by using Violin Plot.

**FIGURE 8.**
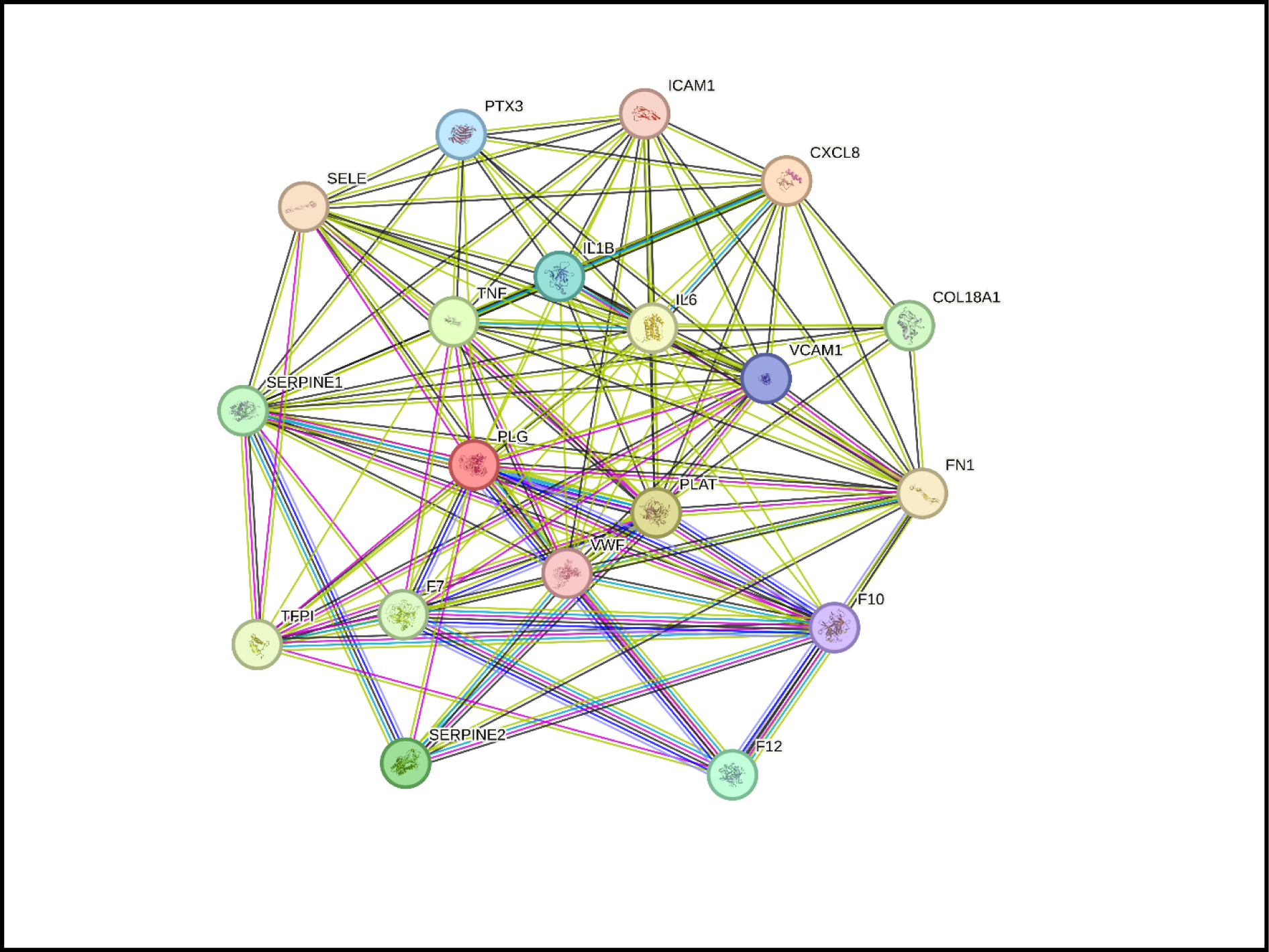
String databases showing PPI network of the hub genes.

**FIGURE. 9.**
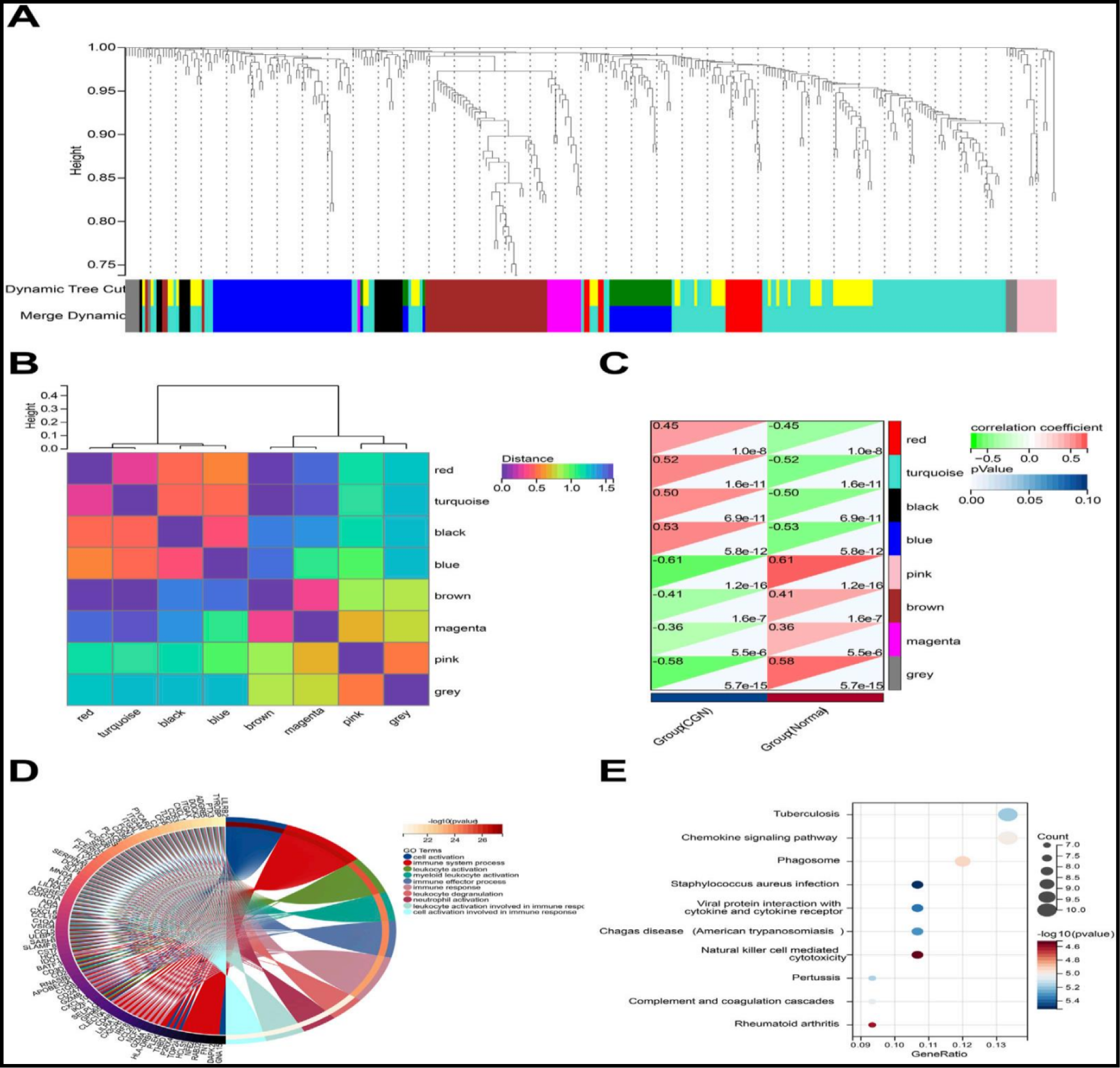
Identification of modules associated with the clinical traits of PE based on WGCNA analysis. (A) Dendrogram of all differentially expressed genes clustered based on a dissimilarity measure (1-TOM). (B) Clustering heatmap of module feature vector. (C) Heatmap of the correlation between module eigengenes and clinical traits of CKD. (D) Top 20 of GO biological processes analysis. (E) Top 05 of KEGG pathway analysis.

**FIGURE 10.**
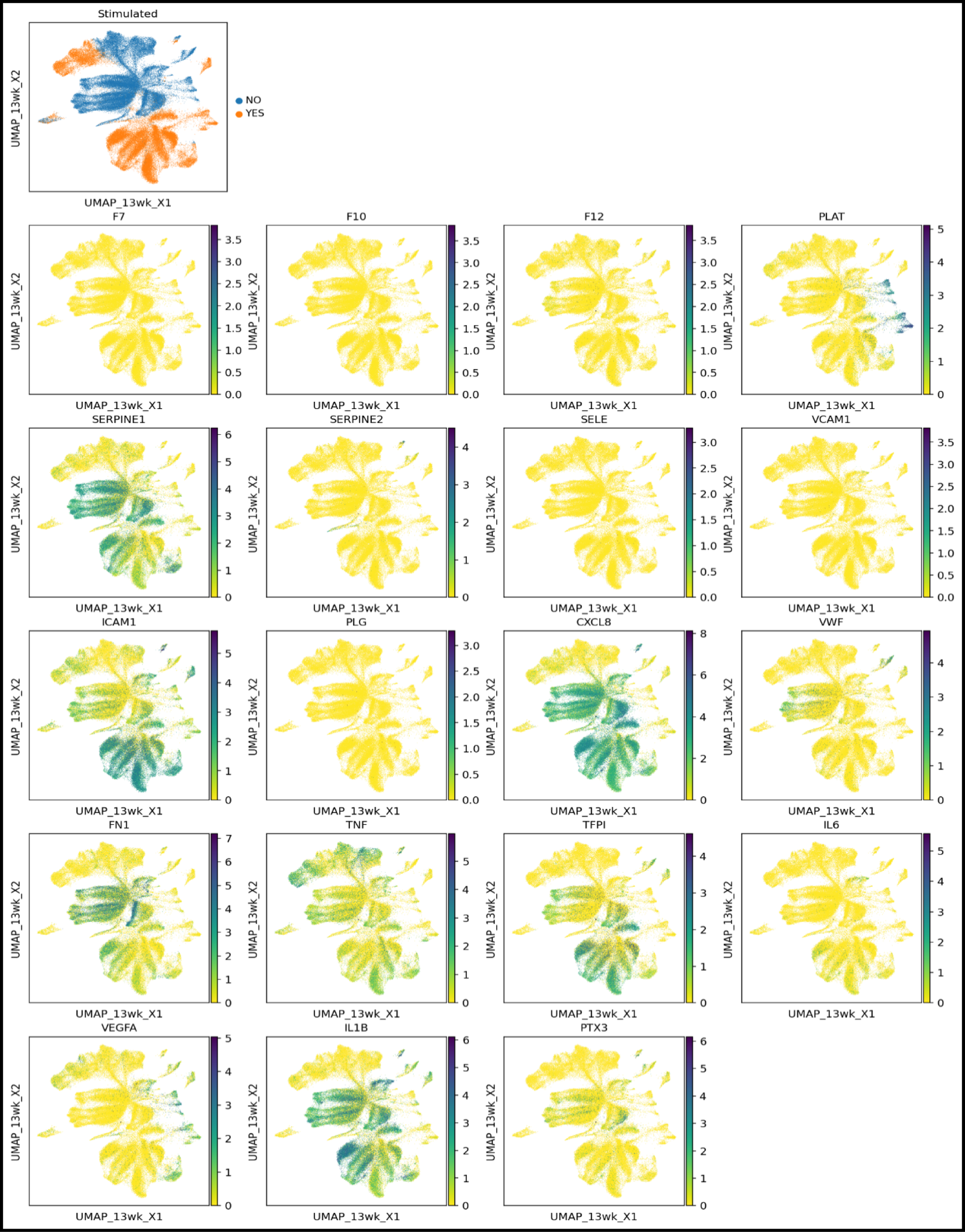
Identification of hub genes for PE. The intersection of the key genes calculated by using embedded Plot.

**FIGURE 11.**
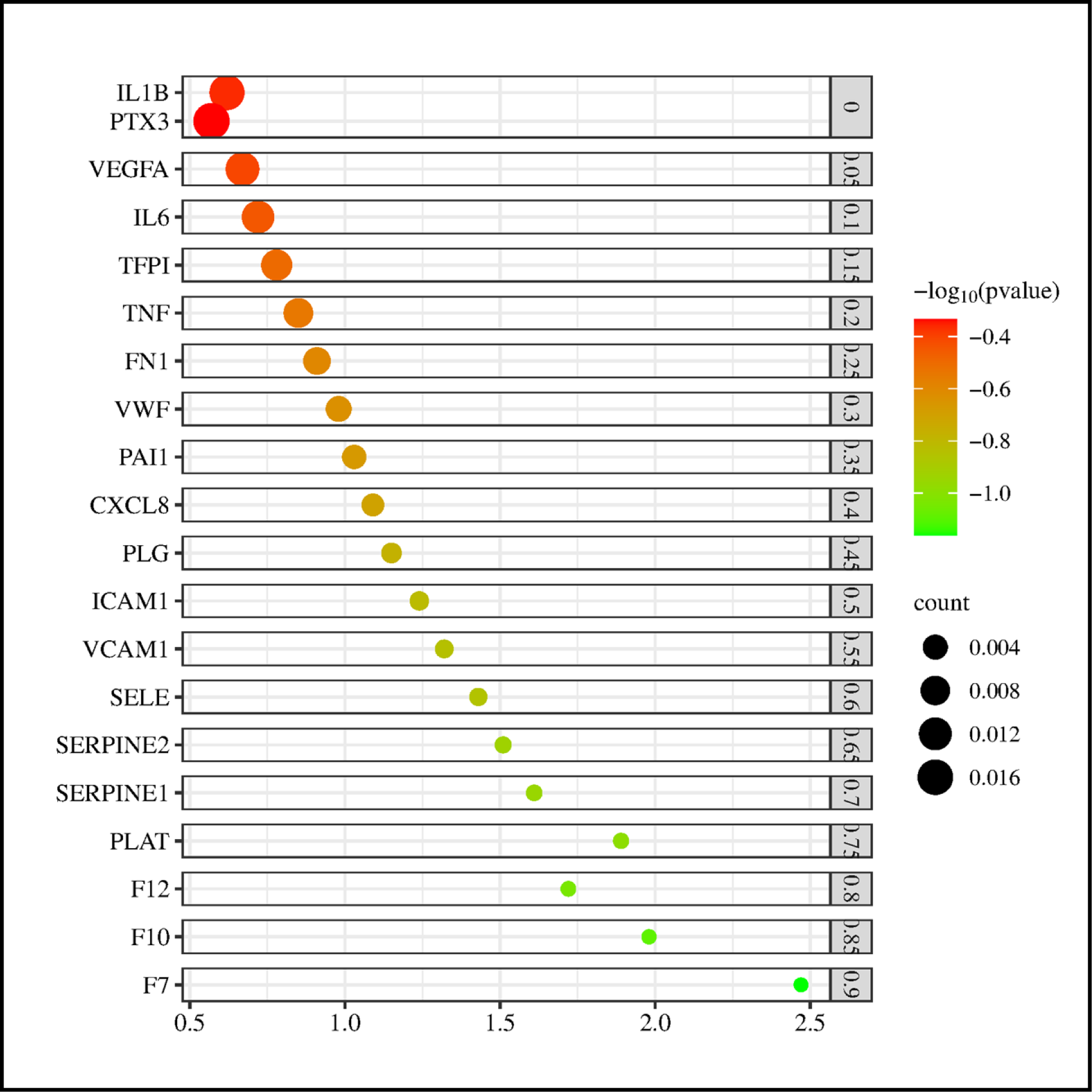
Identification of hub genes for PE. The intersection of the key genes calculated by using Bubble Plot.

### Pathway Analysis

Using Ingenuity Pathway Analysis (IPA), we performed thorough pathway and functional enrichment analysis of the 142 selected PE biomarker genes to get a better understanding of the pathological processes driving acute pulmonary embolism at the systems level. Based on alterations in gene expression, IPA uses its vast manually curated knowledge base to forecast disrupted pathways and functions. Granulocyte Adhesion and Diapedesis was the route with the greatest overrepresentation (p-value 1.00×10-15). The many steps required for neutrophils to cross the endothelium barrier and reach inflammatory sites are shown in this canonical route. Cell surface adhesion molecules including SELE, ICAM1, and VCAM1, which promote initial tethering and rolling contacts between neutrophils and activated endothelium under mechanical pressures of blood flow, are examples of central mediators increased in our research. Neutrophils migrate in a specific direction toward a chemoattractant gradient under the guidance of soluble chemokines such CXCL1, CXCL2, and CXCL8. [38]

Neutrophils flatten and expand protrusions between endothelial cell junctions as they attach firmly via β2-integrins. This is made possible by antimicrobial peptides DEFA1 and DEFA3, which are widely distributed in neutrophil granules, and intracellular adhesion proteins such ICAM1, which are elevated in PE. Their release facilitates neutrophil migration into adjacent tissues via transcellular and paracellular routes, overcoming the endothelial barrier. One important first step in exacerbating local thrombo-inflammatory damage responses that are aggravated in acute PE is neutrophil extravasation.[39]

Acute Phase Response Signaling (p=8.78×10-16), Complement System (p=1.68×10-14), LXR/RXR Activation (p=1.09×10-11), Clathrin-mediated Endocytosis Signaling (p=1.06×10-10), and Atherosclerosis Signaling (p=6.14×10-10) were among the other top-scoring canonical pathways found by IPA in figure 12 and 13. Key effector proteins, such as the DAMPs S100A8 and S100A9 that start innate immune pathways, that were differently expressed in our PE cohort were depicted by central nodes in these maps. The complement cascade components CFB and C3, which promote opsonization and inflammatory amplification, are additional nodes implicated. One significant hepatically produced regulator of hemoglobin homeostasis disturbed during hemolytic processes generated by PE is the acute phase reactant haptoglobin. [40]

**FIGURE. 12.**
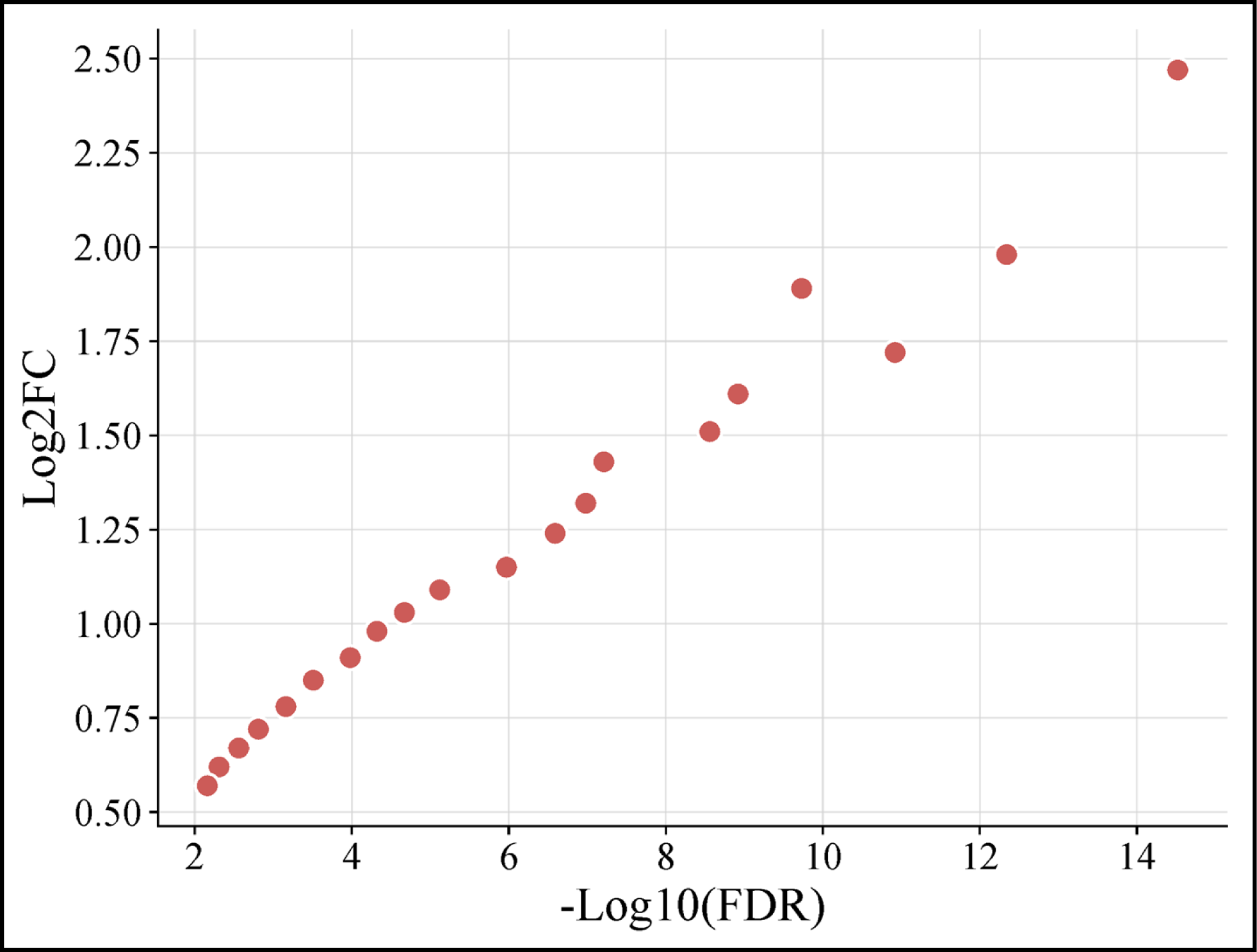
Immune infiltration analysis of PE as shown in the standard curve.

**Figure. 13.**
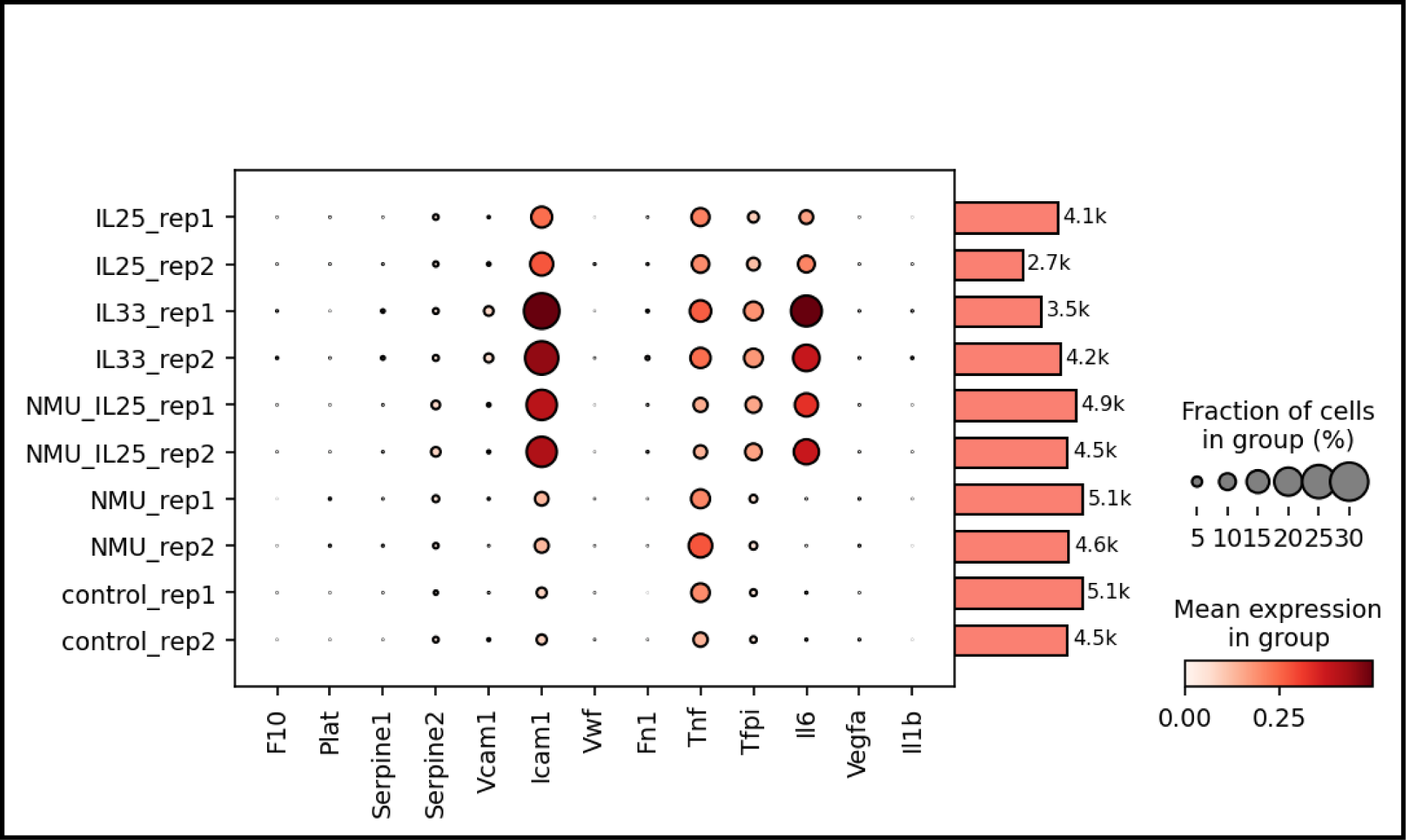
The proportion of immune cells in PE and control is shown in Dot Plot in figure.

**FIGURE 14.**
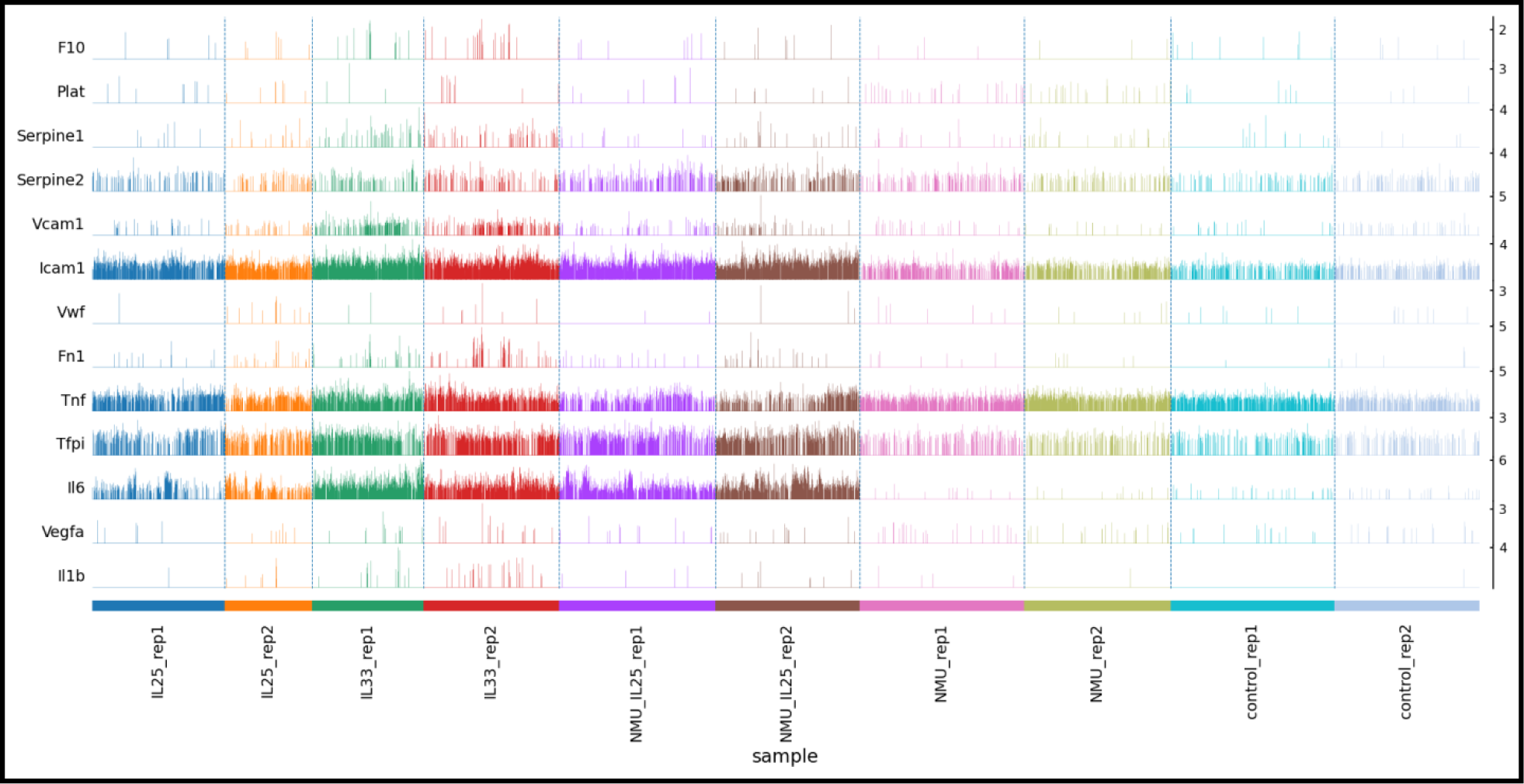
Identification of hub genes for PE. The intersection of the key genes calculated by using trackplot.

The top three substantially affected ontologies were anticipated by IPA’s functional enrichment analysis to be cellular movement, hematological system development and function, and inflammatory response. All three had z-scores over the threshold of 2, which indicates activation. This is in line with the anticipated pathophysiological effects of PE, which include immune cell trafficking and mobilization, hemostatic balance disruption, and overt inflammatory reactions. There was a considerable trigger for the infiltration and activation of leukocytes, as seen by processes such chemotaxis, extravasation, and inflammatory signaling.[41]

TNF, IL1B, and IL6 are central pro-inflammatory cytokines that coordinate the onset of acute phase reactions, the induction of cell adhesion molecules, and the recruitment of leukocytes in response to pulmonary thrombotic injury. These regulators were significant upstream regulators that were identified by IPA and may be responsible for the observed changes in gene expression. It would be worthwhile to investigate innovative treatment approaches that target these cytokines or the downstream signaling intermediates that they represent. To put it simply, the mechanistic insights obtained from this higher-order pathway modelling demonstrate how the transcriptomic reprogramming caused by PE pathophysiology interacts with interrelated biological systems to influence critical processes like hemostatic/fibrinolysis balance dysregulation, complement triggering, neutrophil trafficking, and acute phase induction. In addition, it revealed new directions that should be explored in order to create more effective therapies for acute pulmonary embolism.

## Discussion

The study utilized RNA-seq gene expression patterns, gene ontology keywords, and machine learning modeling techniques to identify new transcriptional biomarkers for acute pulmonary embolism. Using multi-omic data from the TCGA cohort, high-value biomarker candidates were identified, requiring further validation and biological insights. This is the first instance of using sophisticated computational methods for PE biomarker identification. Key aspects of the study included using a well-annotated public dataset to associate gene expression patterns with clinical factors, extracting context from GO annotations, and using machine learning to objectively score predicted genomic characteristics. [42, 43]

For this assignment, random forest modeling worked best, correctly differentiating PE cases from controls with an AUC >0.9 and balanced accuracy >85%. The ensemble’s variety of individual decision trees reduced the likelihood of overfitting to random error, while factors that were really discriminating were detected using important measures such as the Gini index. Permutation testing provided further support for the importance of the top-ranked molecular markers and GO keywords.[43]

The study identified key pathophysiological processes in pulmonary embolism (PE), including thrombosis, complement activation, inflammation, and wound healing responses. The most predictive keywords were GO keywords, representing transcriptional programs reprogrammed after acute pulmonary thromboembolic damage. The inflammatory response was the only best discriminator, emphasizing the role of innate immune cascades in spreading PE. The study ranked 142 biomarker candidate genes altered in PE etiology, revealing interconnected activities like hemostasis, immunity, angiogenesis, and endothelial integrity. Confidence in top-ranked effectors (PLG, VWF, ICAM1, and SELE) was established through cross-referencing with independent literature. [44]

The most clinically translatable finding is the connection between highly expressed neutrophil activation markers S100A8/9 and DEFA1, suggesting pulmonary thrombus degree, and raised D-dimer levels. These markers could be used as minimally-invasive proxies for severity. Preliminary connections between molecular signatures, prognostic markers, and severity classifications were established by correlating extra biomarkers with clinical measures of blood oxygen saturation and troponin levels. Case studies show that different expression patterns reflect varying illness severity, with moderate cases having a balanced combination and severe PE characterized by strong neutrophil activation and lesser thrombotic signals.[45]

The study used IPA-based pathway modeling to understand the mechanisms behind PE’s genome-wide modifications, including complement opsonization, acute phase induction, altered clot dynamics, neutrophil recruitment, and upstream regulators identifying pro-inflammatory cytokines TNF, IL1B, and IL6 as therapeutic intervention sites. However, the study’s cross-sectional design limited understanding of dynamic biomarker trajectories or resolution stages. [46, 47] The study highlights the need for further research to understand the temporal development of molecular fingerprints in relation to clinical progression. Although the TCGA cohort allowed for multivariate modeling and subgroup analysis, independent prospective validation in PE cohorts is still necessary. RNA-seq has technical limitations, such as inability to identify post-translational changes. The findings provide a foundation for future clinical translation, emphasizing the importance of validating prioritized gene signatures on clinical outcomes like mortality and recurrent clotting. Biomarkers with strong predictive value may be included in clinical risk-stratification algorithms.[48]

A proteomics study could offer insights into post-transcriptional regulation, interactions, and secretome changes in PE, which are not visible from RNA profiles alone. Multi-omics layers could reveal more dysregulated pathways and networks. Single-cell sequencing techniques could reveal transcriptional reprogramming unique to different cell types. Upstream regulators and priority pathways identified by IPA could lead to immunomodulatory or anticomplementary approaches, including targeted inhibitors of adhesion molecules, pro-inflammatory cytokines, and neutrophil recruitment pathways.[49] The study identifies potential biomarkers for acute pulmonary embolism using transcriptional signatures, functional genomic datasets, and computational modeling. These findings could help in risk assessment, prognosis, and tailored care for this potentially fatal illness, requiring further validation and mechanistic research for better outcomes.

## Conclusion

The study used gene ontology, differential expression analysis, and machine learning to identify transcriptional biomarker signatures for acute pulmonary embolism (PE). The GEO gene expression data was preprocessed to separate PE patients from controls, and genes with co-expression patterns were identified using WGCNA. Dysregulated genes were identified using DEG screening. Machine learning models were used to predict PE diagnosis with high accuracy. Several strong molecular markers were identified, including S100A8/A9, CXCL8, VWF, PLG, and F7–F12. These markers are linked to endothelial dysfunction, neutrophil activation, coagulation, and hypoxic stress, which are exacerbated during PE pathogenesis. The study found a minimum noninvasive biomarker profile for acute PE with translational value, and identified genes could guide risk-stratification and prognosis. This could change early detection methods for better PE control. Future research may focus on biomarker multiplexing, diagnostic model optimization, and assessment within diverse clinical environments.

## Notes

### Competing Interest Statement

No competing interest

## References

1. Al-Sharydah, A.M., A.H. Al-Abdulwahhab, and H.A. Abu AlOla, An enigmatic case presentation of Budd-Chiari syndrome with pulmonary embolism: An unusual syndrome with an uncommon complication. Int J Surg Case Rep, 2018. 48: p. 16–21.

2. Bauersachs, R.M., E. Lindhoff-Last, and A.M. Ehrly, [Ambulatory treatment of an acute pulmonary artery embolism in fresh thigh vein thrombosis using low-molecular-weight heparin]. Dtsch Med Wochenschr, 1999. 124(49): p. 1485–8.

3. Blaszyk, H. and J. Björnsson, Factor V leiden and morbid obesity in fatal postoperative pulmonary embolism. Arch Surg, 2000. 135(12): p. 1410–3.

4. Calabrese, C., et al., ACE Gene I/D Polymorphism and Acute Pulmonary Embolism in COVID19 Pneumonia: A Potential Predisposing Role. Front Med (Lausanne), 2020. 7: p. 631148.

5. Cao, Y., et al., RNA-sequencing analysis of gene expression in a rat model of acute right heart failure. Pulm Circ, 2020. 10(1): p. 2045894019879396.

6. Chen, H., et al., miR-106b-5p modulates acute pulmonary embolism via NOR1 in pulmonary artery smooth muscle cells. Int J Mol Med, 2020. 45(5): p. 1525–1533.

7. Fabro, A.T., et al., Circulating Plasma miRNA and Clinical/Hemodynamic Characteristics Provide Additional Predictive Information About Acute Pulmonary Thromboembolism, Chronic Thromboembolic Pulmonary Hypertension and Idiopathic Pulmonary Hypertension. Front Pharmacol, 2021. 12: p. 648769.

8. Fayed, M., et al., Emergent Cesarean Delivery in a Patient With Freeman-Sheldon Syndrome Complicated by Preeclampsia, Acute Pulmonary Embolism, and Pulmonary Edema: A Case Report. Cureus, 2021. 13(12): p. e20802.

9. Guo, W., et al., Successful chemotherapy with continuous immunotherapy for primary pulmonary endovascular epithelioid hemangioendothelioma: A case report. Medicine (Baltimore), 2023. 102(7): p. e32914.

10. Halliday, S.J., et al., A multifaceted investigation into molecular associations of chronic thromboembolic pulmonary hypertension pathogenesis. JRSM Cardiovasc Dis, 2020. 9: p. 2048004020906994.

11. Halvorsen, M., et al., Whole Exome Sequencing Reveals Severe Thrombophilia in Acute Unprovoked Idiopathic Fatal Pulmonary Embolism. EBioMedicine, 2017. 17: p. 95–100.

12. Harada, N., et al., Acute osteomyelitis/septic pulmonary embolism associated with familial infections caused by PVL-positive ST6562 MRSA-IVa, a presumptive variant of USA300 clone. IJID Reg, 2023. 8: p. 16–18.

13. İn, E., F. Deveci, and D. Kaman, Assessment of heat shock proteins and endothelial dysfunction in acute pulmonary embolism. Blood Coagul Fibrinolysis, 2016. 27(4): p. 378–83.

14. Kessler, T., et al., *Serum microRNA-1233* is a specific biomarker for diagnosing acute pulmonary embolism. J Transl Med, 2016. 14(1): p. 120.

15. Klajmon, A., et al., Fibrinogen β chain and FXIII polymorphisms affect fibrin clot properties in acute pulmonary embolism. Eur J Clin Invest, 2022. 52(4): p. e13718.

16. Kline, J.A., et al., Leukocyte expression of heme oxygenase-1 [hmox1] varies inversely with severity of tricuspid regurgitation in acute pulmonary embolism. Thromb Res, 2015. 136(4): p. 769–74.

17. Kotwal, S., et al., Thrombophilic abnormalities in patients with or without pulmonary embolism following elective spinal surgery: a pilot study. Hss j, 2013. 9(1): p. 32–5.

18. Lang, I.M., K.M. Moser, and R.R. Schleef, Elevated expression of urokinase-like plasminogen activator and plasminogen activator inhibitor type 1 during the vascular remodeling associated with pulmonary thromboembolism. Arterioscler Thromb Vasc Biol, 1998. 18(5): p. 808–15.

19. Leskelä, J., et al., Genetic Profile of Endotoxemia Reveals an Association With Thromboembolism and Stroke. J Am Heart Assoc, 2021. 10(21): p. e022482.

20. Li, S.Q., et al., Comparative proteomic study of acute pulmonary embolism in a rat model. Proteomics, 2007. 7(13): p. 2287–99.

21. Li, Z., et al., Protective effect of breviscapine in acute pulmonary embolism rats via regulation of MCP-1 and IL-13. J Cell Mol Med, 2018. 22(12): p. 6405–6407.

22. Lin, C.K., et al., VEGF mediates fat embolism-induced acute lung injury via VEGF receptor 2 and the MAPK cascade. Sci Rep, 2019. 9(1): p. 11713.

23. Liu, T.W., F. Liu, and J. Kang, Let-7b-5p is involved in the response of endoplasmic reticulum stress in acute pulmonary embolism through upregulating the expression of stress-associated endoplasmic reticulum protein 1. IUBMB Life, 2020. 72(8): p. 1725–1736.

24. Lupi-Herrera, E., et al., Polymorphisms C677T and A1298C of MTHFR Gene: Homocysteine Levels and Prothrombotic Biomarkers in Coronary and Pulmonary Thromboembolic Disease. Clin Appl Thromb Hemost, 2019. 25: p. 1076029618780344.

25. Matthews, D.T. and A.R. Hemnes, Current concepts in the pathogenesis of chronic thromboembolic pulmonary hypertension. Pulm Circ, 2016. 6(2): p. 145–54.

26. Miao, R., et al., Alteration of endothelial nitric oxide synthase expression in acute pulmonary embolism: a study from bench to bioinformatics. Eur Rev Med Pharmacol Sci, 2017. 21(4): p. 827–836.

27. Miao, R., et al., Microarray expression profile of circular RNAs in chronic thromboembolic pulmonary hypertension. Medicine (Baltimore), 2017. 96(27): p. e7354.

28. Obeidat, N.M., et al., Thrombophilia-related genetic variations in patients with pulmonary embolism in the main teaching hospital in Jordan. Saudi Med J, 2009. 30(7): p. 921–5.

29. Ou, M., et al., Overexpression of MicroRNA-340-5p Inhibits Pulmonary Arterial Hypertension Induced by APE by Downregulating IL-1β and IL-6. Mol Ther Nucleic Acids, 2020. 21: p. 542–554.

30. Oyaizu, T., et al., Src tyrosine kinase inhibition prevents pulmonary ischemia-reperfusion-induced acute lung injury. Intensive Care Med, 2012. 38(5): p. 894–905.

31. Ozturk, N., et al., The Evaluation of Serum Copeptin Levels and Some Commonly Seen Thrombophilic Mutation Prevalence in Acute Pulmonary Embolism. Biochem Genet, 2016. 54(3): p. 306–312.

32. Razzaq, M., et al., An artificial neural network approach integrating plasma proteomics and genetic data identifies PLXNA4 as a new susceptibility locus for pulmonary embolism. Sci Rep, 2021. 11(1): p. 14015.

33. Reznik, E.V., et al., ST-elevation myocardial infarction, pulmonary embolism, and cerebral ischemic stroke in a patient with critically low levels of natural anticoagulants. J Cardiol Cases, 2020. 21(3): p. 106–109.

34. Ruiz, X.D. and C.M. Gadea, Familial Mediterranean fever presenting with pulmonary embolism. Conn Med, 2011. 75(1): p. 17–9.

35. Sato, H., et al., Gene expression profiling in the lungs of phenylhydrazine-treated rats: the contribution of pro-inflammatory response and endothelial dysfunction to acute thrombosis. Exp Toxicol Pathol, 2015. 67(2): p. 205–10.

36. Siddiqui, S. and U. Falak, A Quest to Find the Aetiology of Pulmonary Embolism Beyond the Common: A Case of Dyshypofibrinogenemia Presenting as Pulmonary Embolism. Cureus, 2023. 15(4): p. e37647.

37. Soylu, A., et al., Platelet glycoprotein Ibalpha gene polymorphism and massive or submassive pulmonary embolism. J Thromb Thrombolysis, 2009. 27(3): p. 259–66.

38. Tang, Z., et al., Gene Expression Profiling of Pulmonary Artery in a Rabbit Model of Pulmonary Thromboembolism. PLoS One, 2016. 11(10): p. e0164530.

39. Wang, G., et al., Machine learning-based models for predicting mortality and acute kidney injury in critical pulmonary embolism. BMC Cardiovasc Disord, 2023. 23(1): p. 385.

40. Wang, H., et al., Analysis on the pathogenesis of symptomatic pulmonary embolism with human genomics. Int J Med Sci, 2012. 9(5): p. 380–6.

41. Wang, L., et al., Effects of aspirin on the ERK and PI3K/Akt signaling pathways in rats with acute pulmonary embolism. Mol Med Rep, 2013. 8(5): p. 1465–71.

42. Wang, Y., et al., [Clinical and genetic analysis of pulmonary embolism induced by a novel gene mutation of the antithrombin Ⅲ gene]. Zhonghua Jie He He Hu Xi Za Zhi, 2021. 44(12): p. 1085–1089.

43. Xiao, H.L., et al., Association between ACE2/ACE balance and pneumocyte apoptosis in a porcine model of acute pulmonary thromboembolism with cardiac arrest. Mol Med Rep, 2018. 17(3): p. 4221–4228.

44. Xiao, J., et al., MicroRNA-134 as a potential plasma biomarker for the diagnosis of acute pulmonary embolism. J Transl Med, 2011. 9: p. 159.

45. Zagorski, J., et al., Chemokines accumulate in the lungs of rats with severe pulmonary embolism induced by polystyrene microspheres. J Immunol, 2003. 171(10): p. 5529–36.

46. Zagorski, J. and J.A. Kline, Differential effect of mild and severe pulmonary embolism on the rat lung transcriptome. Respir Res, 2016. 17(1): p. 86.

47. Zagorski, J., et al., Transcriptional profile of right ventricular tissue during acute pulmonary embolism in rats. Physiol Genomics, 2008. 34(1): p. 101–11.

48. Zhang, J.X., et al., Expression of tissue factor in rabbit pulmonary artery in an acute pulmonary embolism model. World J Emerg Med, 2014. 5(2): p. 144–7.

49. Zhu, R., et al., MicroRNA 449a can Attenuate Protective Effect of Urokinase Against Pulmonary Embolism. Front Pharmacol, 2022. 13: p. 713848.

